# Beclin1-Deficient Adipocytes Promote Tumor Progression by YAP/TAZ-dependent Adipocyte Transformation

**DOI:** 10.1101/2023.05.25.542253

**Authors:** Yaechan Song, Heeju Na, Seung Eon Lee, Jihyun Moon, Tae Wook Nam, You Min Kim, Yul Ji, Young Jin, Jae Hyung Park, Seok Chan Cho, Daehee Hwang, Sang-Jun Ha, Hyun Woo Park, Jae Bum Kim, Han-Woong Lee

## Abstract

Adipocytes are crucial components of the tumor microenvironment (TME) that play a prominent role in supporting tumor growth. However, the characteristics of cancer-associated adipocytes (CAAs) that contribute to the pro-tumorigenic niche remain to be fully established. Here, we used adipocyte-specific *Beclin1 KO* (BaKO) mice to investigate the role of maladaptive adipocytes in promoting tumor progression. BECN1-deficient adipocytes exhibited downregulation of adipogenic markers and activation of YAP/TAZ signaling, similar to the traits observed in CAAs. Thus, we generated adipocyte-specific *Becn1/Yap1/Taz KO* mice, which exhibit markedly restored phenotypes in adipose tissue, resulting in tumor regression compared to that in BaKO. Further, we observed dysregulation of the BECN1-YAP/TAZ axis in the adipose tissue of mice fed a high-fat diet (HFD). Treatment with the YAP/TAZ inhibitor, verteporfin, suppressed tumor progression in BaKO and HFD-fed mice, highlighting its efficacy against mice with metabolic dysregulation. Our findings provide insights into CAA formation and its significance in determining malignant TME, thereby suggesting a potential dual therapeutic strategy simultaneously targeting adipocyte homeostasis and cancer growth.

## Introduction

The study of tumor progression goes beyond cancer cells to encompass the complex ecosystem surrounding them. This ecosystem is the tumor microenvironment (TME), comprising a variety of cell types, including immune, epithelial, and endothelial cells, fibroblasts, and adipocytes. Among the TME compartments, adipocytes make up a considerable portion of breast, colorectal, and endometrial stroma (Pallegar & Christian, 2020; Rybinska *et al*, 2021). Adipocytes located at the tumor protruding region undergo dynamic morphological and functional transformations. These adipocyte populations, known as cancer-associated adipocytes (CAAs), exhibit dedifferentiated phenotypes and fibroblast-like morphologies (Bochet *et al*, 2013; Yao & He, 2021), undergoing metabolic adaptation in a nutrient-restrictive environment (Hoy *et al*, 2017).

CAAs prominently activate lipolysis to release lipid metabolites, such as free fatty acids (FFAs), which cancer cells utilize to fuel fatty acid oxidation (FAO) for rapid progression (Balaban *et al*, 2017; Martin-Perez *et al*, 2022). Upon transformation, adipocytes secrete adipokines and signaling molecules to provide a pro-tumorigenic environment (Gnerlich *et al*, 2013). However, there is a lack of literature characterizing CAAs, especially how they are generated and what key elements contribute to cancer progression.

Adipose tissue in obese individuals is subjected to hypoxic conditions due to the increased number and size of adipocytes. These adipocytes secrete pro-inflammatory factors, such as TNFα, interleukin-6 (IL-6), monocyte chemoattracted protein-1 (MCP1), and CXC motif chemokine ligand 12 (CXCL12), which cause local and systemic inflammation (Ellulu *et al*, 2017). Adipose tissue dysfunction alters endocrine signals, exacerbating diverse illnesses, such as diabetes, cardiovascular disease, hepatic steatosis, and cancer. The state of adipocytes can be a strong determinant of disease progression, as it reflects the cellular state of energy metabolism, hormone regulation, and immune function (Kawai *et al*, 2021). However, only a few studies have attempted to devise a therapeutic strategy to target adipocyte transformation.

*Beclin1* (*BECN1*) is an essential autophagy-related gene that initiates autophagy by activating the class 3 phosphoinositide 3-kinase (PI3K) complex through ULK1-mediated BECN1 phosphorylation (Kang *et al*, 2011). However, recent studies have highlighted various autophagy-independent functions of BECN1 (Galluzzi & Green, 2019). BECN1 is involved in cellular processes that require membrane remodeling, such as vacuolar protein sorting, cytokinesis, endocytosis, and phagocytosis (Galluzzi & Green, 2019). According to the clinical data, BECN1 has been suggested to have tumor-suppressive functions since monoallelic *Becn1* deletion has been found in ∼30% of breast cancer and ∼77% of ovarian cancer patients (Liang *et al*, 1999; Qu *et al*, 2003; Yue *et al*, 2003). Hu *et al*. also reported that downregulation of BECN1 expression in patients with colorectal cancer led to the suppression of cancer metastasis in mice through inhibition of STAT3 phosphorylation via Janus kinase 2 (JAK2) (Hu *et al*, 2020). While the intrinsic tumor function of BECN1 has been well studied, much remains to be unveiled in the context of the TME. In this study, we aimed to investigate the function of BECN1 in the adipocyte-TME, given the crucial role adipocytes play in promoting tumor growth and their unique metabolic adaptations in response to nutrient restriction.

In our previous study, we generated adipocyte-specific *Becn1 KO* mice (BaKO) to investigate the role of BECN1 in mature adipocytes. We reported that BECN1-deficient adipocytes experienced a downregulation of Perilipin1 (PLIN1) expression and loss of lipid contents. BECN1 depletion also induced endoplasmic reticulum (ER) stress, leading to adipocyte death through enhanced apoptotic signals (Jin *et al*, 2021). In this study, we investigate the clinical implication of BECN1 loss in adipocytes, since it could facilitate a pro-tumorigenic niche in adipose-rich tumor microenvironment. We show that the loss of BECN1 leads to adipocyte transformation, which displayed characteristics similar to the peritumoral and cancer-co-cultured adipocytes through activation of YAP/TAZ signaling and inflammatory response. To abrogate the impact of YAP/TAZ-mediated adipocyte transformation, we generated adipocyte-specific *Becn1/Yap1/Taz* triple-KO mice (BYTaKO). Here, inflammatory and pro-tumorigenic adipokines were curtailed, leading to tumor regression in BYTaKO. These findings highlight the significance of BECN1-mediated YAP/TAZ regulation in the adipocyte-TME and suggest that inhibition of YAP/TAZ to retain adipocyte quality may provide a novel therapeutic approach to target TME compartments.

## Results

### BECN1-deficient adipocytes exhibit dedifferentiated phenotypes, as indicated by YAP and β- catenin

Autophagy, a cellular process of degradation and recycling of unwanted cellular components, is an essential process for adipogenesis (Tao *et al*, 2019; Zhang *et al*, 2013). Since adipocyte differentiation involves dynamic changes in gene expression, we aimed to investigate how the autophagy-related protein, BECN1, is regulated during this process. We observed that BECN1 expression increases as adipocyte differentiation progresses. In contrast, signaling molecules for cellular plasticity and proliferation, such as YAP/TAZ and β-catenin, were downregulated during adipocyte differentiation (Fig. 1A and B). As we further explored the role of YAP/TAZ in adipocyte maturity, our IF analysis showed that the mature adipocytes expressing high PLIN1 levels have low nuclear YAP/TAZ levels, whereas PLIN1-negative preadipocytes contain noticeably higher levels of nuclear YAP/TAZ. Thus, YAP/TAZ localization serves as an indicator of adipocyte maturity (Fig EV1A).

**Figure 1.**
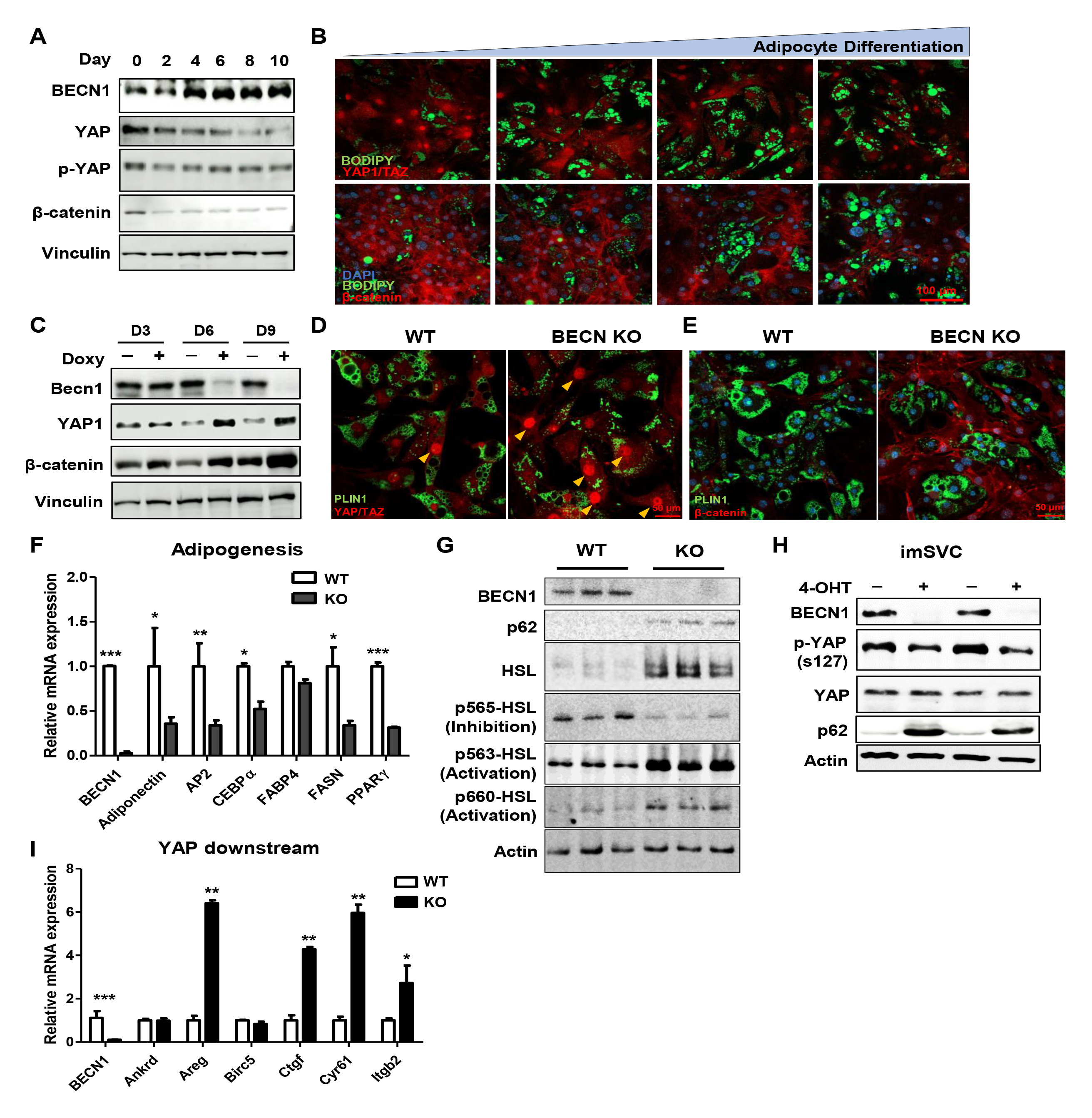
BECN1-deficient adipocytes lose maturity, as indicated by YAP and β-catenin activity. A Western blotting analysis of protein lysates isolated from differentiating adipocytes over the time course. B Representative IF images throughout adipocyte differentiation. Adipocytes were stained with YAP/TAZ and β-catenin (red), BODIPY (green), and DAPI (blue). C Western blotting analysis of protein lysates isolated from doxy-inducible *Becn1 KO* cell lines. Fully differentiated adipocytes with and without doxycycline (Doxy) treatment for the time indicated. D Representative IF images of mature adipocytes stained with YAP/TAZ (D) and β-catenin (E). Arrowheads indicate nuclear YAP/TAZ. F Relative mRNA expression of adipogenic genes from conditional *Becn1 KO* adipocytes (imSVC), with and without 4-OHT (n = 3) treatment. G, H Western blotting analysis of protein lysates isolated from imSVCs with and without 4-OHT treatment. I Relative mRNA expression of YAP/TAZ downstream genes from imSVC with and without 4-OHT treatment (n=3). Data information: Two-tailed unpaired Students *t-*test (F and I). Data are shown as mean ± SEM; \**p* ≤ 0.05, \*\**p* ≤ 0.01, \*\*\**p* ≤ 0.001.

To elucidate the physiological response to BECN1 depletion in adipocytes, we established doxycycline-inducible *Becn1 KO* adipocytes. Our results showed that YAP and β-catenin levels were inversely correlated with BECN1 levels (Fig. 1C). IF imaging also revealed enhanced levels of YAP/TAZ and β-catenin in *Becn1 KO* adipocytes, which displayed features of undifferentiated adipocytes (Fig. 1D and E; Fig EV1B-E). To characterize adipocyte transformation, we used conditional *Becn1 KO* adipocytes (imSVC), in which *Becn1* could be deleted by treatment with 4-OHT [method described in (Jin *et al*., 2021)]. BECN1 depletion in imSVCs led to the suppression of adipogenic potential (Fig. 1F) and loss of lipid content due to a marked increase in lipolysis (Fig. 1G; Fig EV1B and F). Consistent with doxycycline-inducible *Becn1 KO* adipocytes, 4-OHT-treated imSVCs exhibited enhanced nuclear translocation of YAP/TAZ and their downstream signaling proteins (Fig. 1H and I). These findings show that BECN1 depletion induces adipocyte transformation, leading to dedifferentiated phenotypes followed by β-catenin expression and YAP/TAZ activity.

### BECN1-deficient adipocytes exhibit similar features to CAAs

When influenced by adjacent tumor cells, CAAs undergo a dynamic transformation to support cancer cell growth (Fig EV1G). CAAs exhibit fibroblast-like phenotypes while losing their mature adipocyte properties (Fig EV1H) (Mukherjee *et al*, 2022; Rybinska *et al*., 2021; Wu *et al*, 2023; Zoico *et al*, 2016). Indeed, when adipocytes were cultivated with breast cancer cells, they expressed remarkably low levels of adipogenic markers, in addition to BECN1 depletion (Fig. 2 A and B). As observed in BECN1- deficient adipocytes, co-cultivated adipocytes expressed enhanced YAP/TAZ activity (Fig. 2C). Another key feature of CAAs is active lipolysis, which releases a substantial amount of lipid content to support tumor growth (Balaban *et al*., 2017; Martin-Perez *et al*., 2022). Similar to CAAs, BECN1- deficient adipocytes have shown persistent activation of lipolysis and release of FFAs into the cultured media (Fig. 1G; Fig EV1F). Collectively, the comparable characteristics between BECN1-deficient adipocytes and *in-vitro* CAAs indicate that BECN1 depletion causes severe adipocyte transformation.

**Figure 2.**
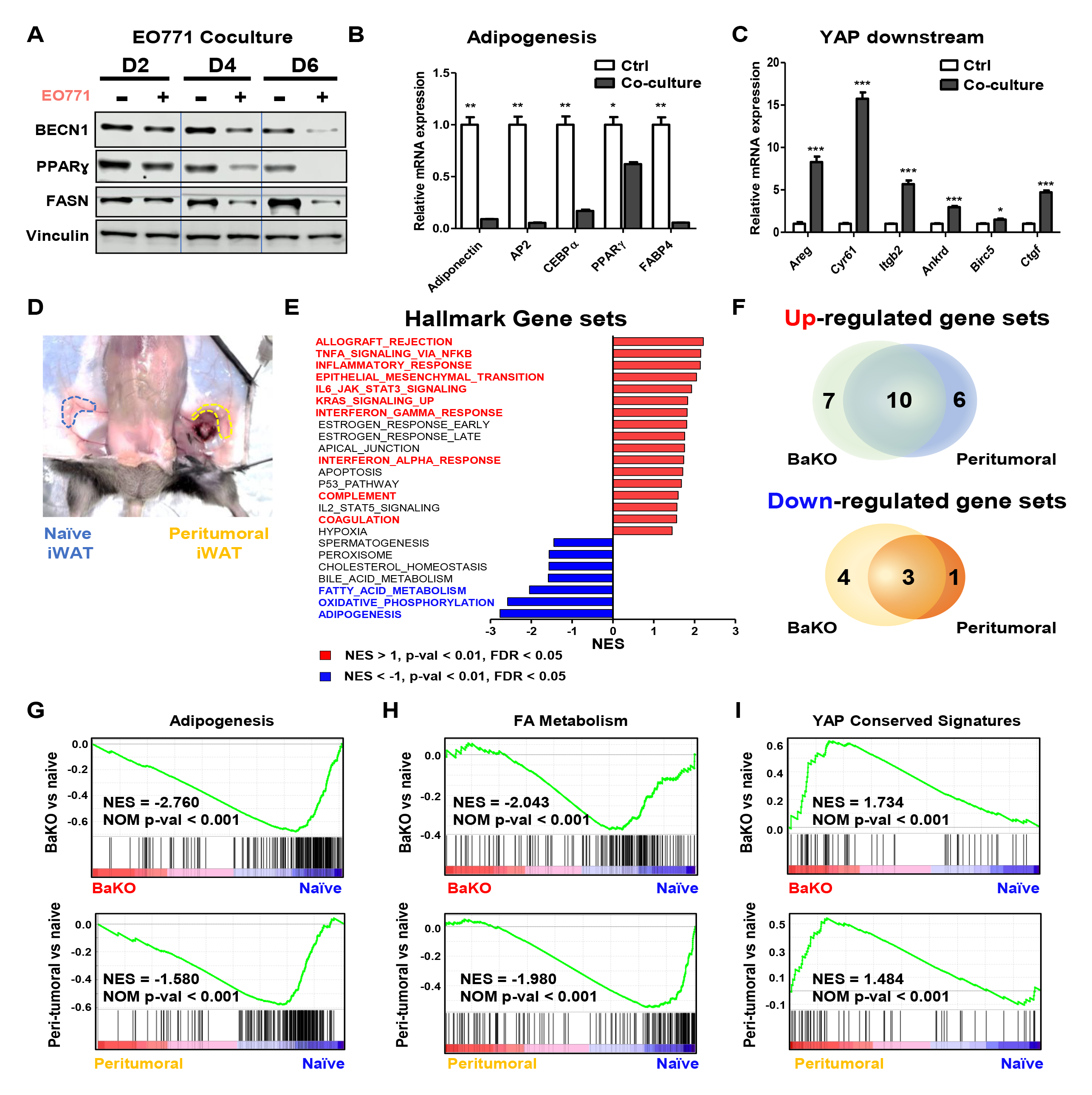
BECN1-deficient adipocytes undergo transformation and exhibit similar features to peritumoral adipocytes. A Western blotting analysis of protein lysates isolated from differentiated 10T1/2 adipocytes co-cultured with EO771 breast cancer cell line over the time course. B Relative mRNA expression of adipogenic genes (B) and YAP/TAZ downstream genes from differentiated 10T1/2 adipocytes co-cultured with EO771 breast cancer cell lines for four days (C). D Image of naïve and peri-tumoral iWAT samples resected from a mouse. E List of differentially expressed GSEA hallmark gene sets of BaKO and naïve adipose tissue ranked by normalized enrichment score (NES). Gene sets that also appear to be differentially expressed in peritumoral WAT are colored in red and blue. F Venn diagram depicting differentially expressed GSEA hallmark gene sets of BaKO and peritumoral WATs compared to naïve adipose tissue; *p* < 0.01, FDR <0.05, NES < −1 or NES > 1 G GSEA plots for adipogenesis (G), FA metabolism (H), and YAP conserved signaling gene sets (I) compared between naïve, peritumoral, and BaKO WAT. Data information: Two-tailed unpaired students *t*-test (B and C). Data are shown as mean ± SEM; \**p* ≤ 0.05, \*\**p* ≤ 0.01, \*\*\**p* ≤ 0.001.

Next, we sought to characterize the CAAs and BECN1-deficient adipocytes *in vivo*. We performed RNA-seq to understand the physiological processes that WATs undergo upon interaction with cancer cells (Fig. 2D-I; Fig EV2A-E). Investigation of the RNA-seq results from naïve, peritumoral, and BaKO WATs further highlighted the resemblance between BECN1-deficient adipocytes and CAAs. We compared the individual genes that were upregulated in BaKO and peritumoral WATs than in naïve WAT (Fig EV2A). The hallmark gene set enrichment analysis (HGSEA) identified that among the 23 upregulated and 8 downregulated gene sets in peritumoral and BaKO WATs, they shared 10 and 3 gene sets to be co-regulated when compared to naïve WAT, respectively (Fig. 2E-F; Fig EV2B).

Zhu *et al*. observed that adipocytes located near cancer cells undergo dedifferentiation under restrictive metabolic conditions, contributing to the generation of a tumor-friendly microenvironment through adipocyte mesenchymal transition (Zhu *et al*, 2022b). Our HGSEA results indicate that epithelial-mesenchymal transition (EMT) was one of the commonly upregulated gene sets in both BaKO and peritumoral WATs (Fig. 2E; Fig EV2B and C). Correspondingly, the gene sets associated with adipogenesis and fatty acid metabolism were co-regulated, supporting the notion of mesenchymal transition of adipocytes (Fig. 2G and H) (Zhu *et al*., 2022b; Zoico *et al*., 2016). We further confirmed the amplified YAP signaling in BaKO, which was also upregulated in peritumoral WAT (Fig. 2I; Fig EV2D-F). Overall, acquisition of peritumoral features by BaKO WAT could create a favorable environment for tumor growth before cancer cell injection.

### BaKO provides a favorable environment for malignant tumor progression

Subsequently, we investigated whether early exposure to dysfunctional adipocytes in the TME could lead to accelerated tumor growth in mice. We employed five syngeneic tumor models and found that colorectal (MC-38) and breast cancer (EO771) cells exhibited more aggressive progression in BaKO (Fig. 3A and B; Appendix Fig S1A and B). However, the progression was not as prominent in melanoma and lung cancer models (Appendix Fig S1C-E).

**Figure 3.**
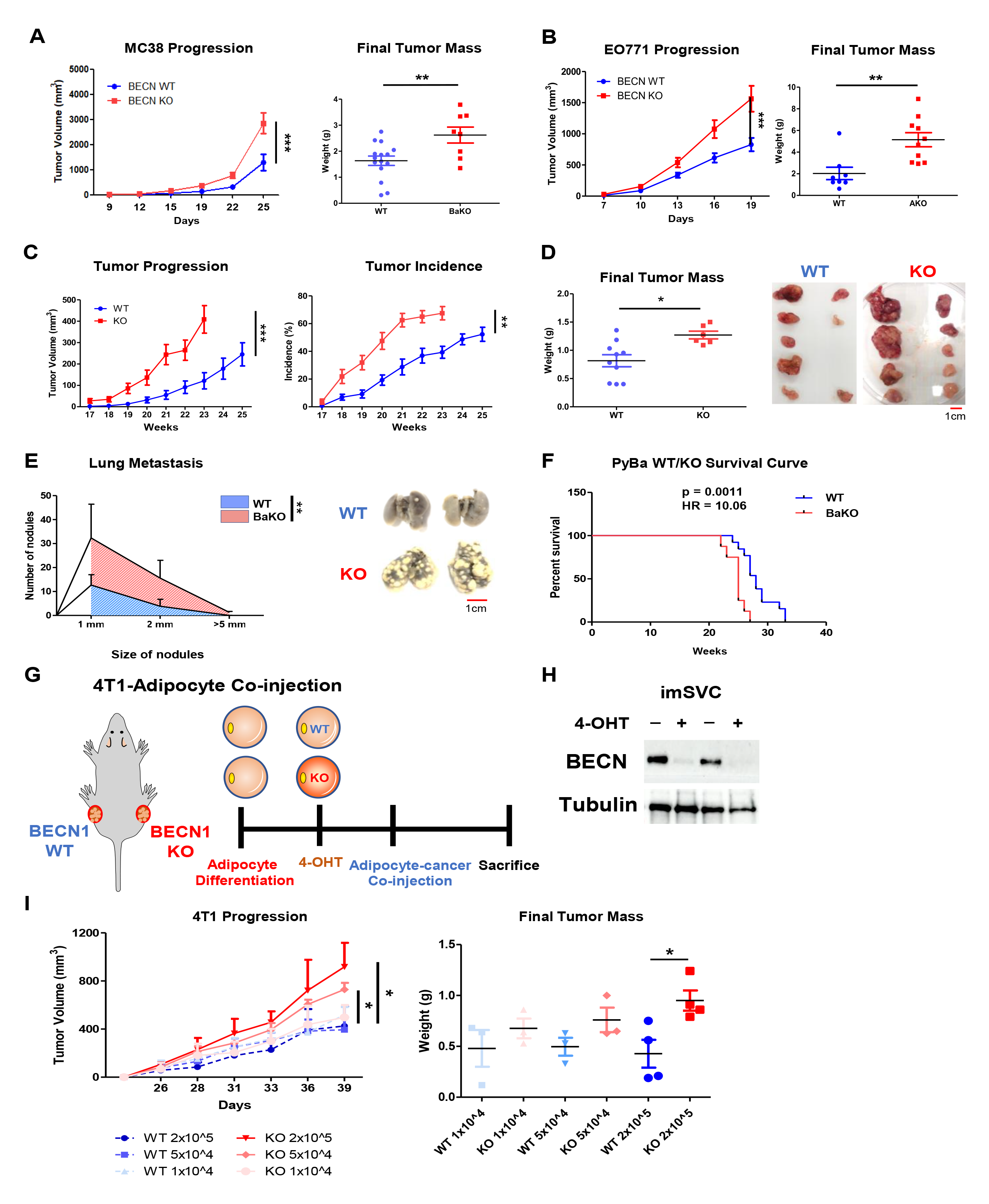
Beclin1 deficient adipocytes promote tumor malignancy in breast and colon cancer models. A Tumor volumes and weights after subcutaneous injection of MC-38 into 6- week-old WT and BaKO mice (WT, n = 9; BaKO, n = 11). B Tumor volumes and weights after mammary fat pad injection of EO771 into 6-week-old WT and BaKO mice (WT, n = 18; BaKO, n = 12). C Average growth kinetics and incidence of breast tumors generated from 10 mammary glands of PyWT and PyBaKO (n = 16). D Average tumor mass isolated from each mammary gland and a representative image of the tumors from PyWT (n = 10) and PyBaKO (n = 6). Mice were sacrificed at 23 weeks of age. E Metastasized tumor nodules counted from the lungs of PyWT (n = 13) and PyBaKO (n = 8) at 23 weeks of age. F Kaplan–Meier plot of survival curve of PyWT and PyBaKO. Log-rank test p-value (p) and HR between PyWT (n = 13) and PyBaKO (n = 8) are indicated. G Schematic timeline of adipocyte-cancer co-injection. H Western blotting analysis of protein lysates isolated from imSVCs before being mixed with cancer cells. I Tumor volumes and weights after subcutaneous injection of 4T1 cells mixed with WT or *Becn1 KO* adipocytes. Mice were sacrificed on the 39^th^ day following co-injection (n= 3,4). Data information: Two-tailed unpaired Student’s *t*-test (A, B, D, and I), ordinary two-way ANOVA (A, B, C, F, and I), and Kaplan–Meier survival curve analysis (e). Data are shown as mean ± SEM; \**p* ≤ 0.05, \*\**p* ≤ 0.01, \*\*\**p* ≤ 0.001.

Furthermore, we utilized a well-established transgenic mouse mammary tumor virus polyoma middle T antigen (MMTV-PyMT) model that spontaneously develops breast cancer (Lin *et al*, 2003). Here, we generated adipocyte-specific *Becn1 KO* conditions during breast cancer development by crossing MMTV-PyMT mice (PyWT) with BaKO (PyBaKO). PyBaKO mice exhibited an increased tumor incidence and progression (Fig. 3C and D), resulting in higher lung metastasis and mortality rates (Fig. 3E and F). These findings, in line with our transcriptomic analysis, underscore the role of BaKO iWAT in providing a tumor-supportive niche for promoting tumor progression.

Notably, in our previous study, we reported that BaKO mice develop lipodystrophy, liver steatosis, and glucose intolerance (Jin *et al*., 2021). These metabolic dysregulations are known to promote tumor progression through systemic impacts (Braun *et al*, 2011; Lega & Lipscombe, 2020), which could have contributed to the rapid tumor growth observed in BaKO mice. To determine the direct effect of BECN1-deficient adipocytes, we co-injected WT or *Becn1 KO* imSVCs with breast cancer cells (4T1) into mice. Prior to mixing with cancer cells, differentiated adipocytes were treated with 4-OHT to deplete BECN1, followed by injection into the mouse flanks (Fig. 3G and H). Interestingly, tumors grew more rapidly as the proportion of *Becn1* KO adipocytes in the mix increased, which was not observed in cancer cells mixed with WT adipocytes (Fig. 3I). These data demonstrate that *Becn1 KO* adipocytes were sufficient to promote tumor growth without the systemic effects observed in BaKO mice.

### BECN1-deficient adipocytes induce inflammatory signals, TNFα and LCN2, to support tumor growth

To elucidate the functional consequences of BECN1 deficiency in adipocytes, we focused our investigation on the considerably upregulated gene sets observed in the transcriptomic analysis of BaKO iWAT. Our results indicated that the acquisition of inflammatory adipose tissue in BaKO was accompanied by the enrichment of gene sets associated with the inflammatory response and TNFα signaling (Fig. 4A and B). Further examination of TNFα downstream signaling revealed its activation in *Becn1 KO* imSVC (Fig. 4C). We also observed that BECN1-deficient adipocytes secrete elevated levels of TNFα into the cultured media, comparable to those seen in normal adipocytes treated with lipopolysaccharide (LPS) (Fig. 4D).

**Figure 4.**
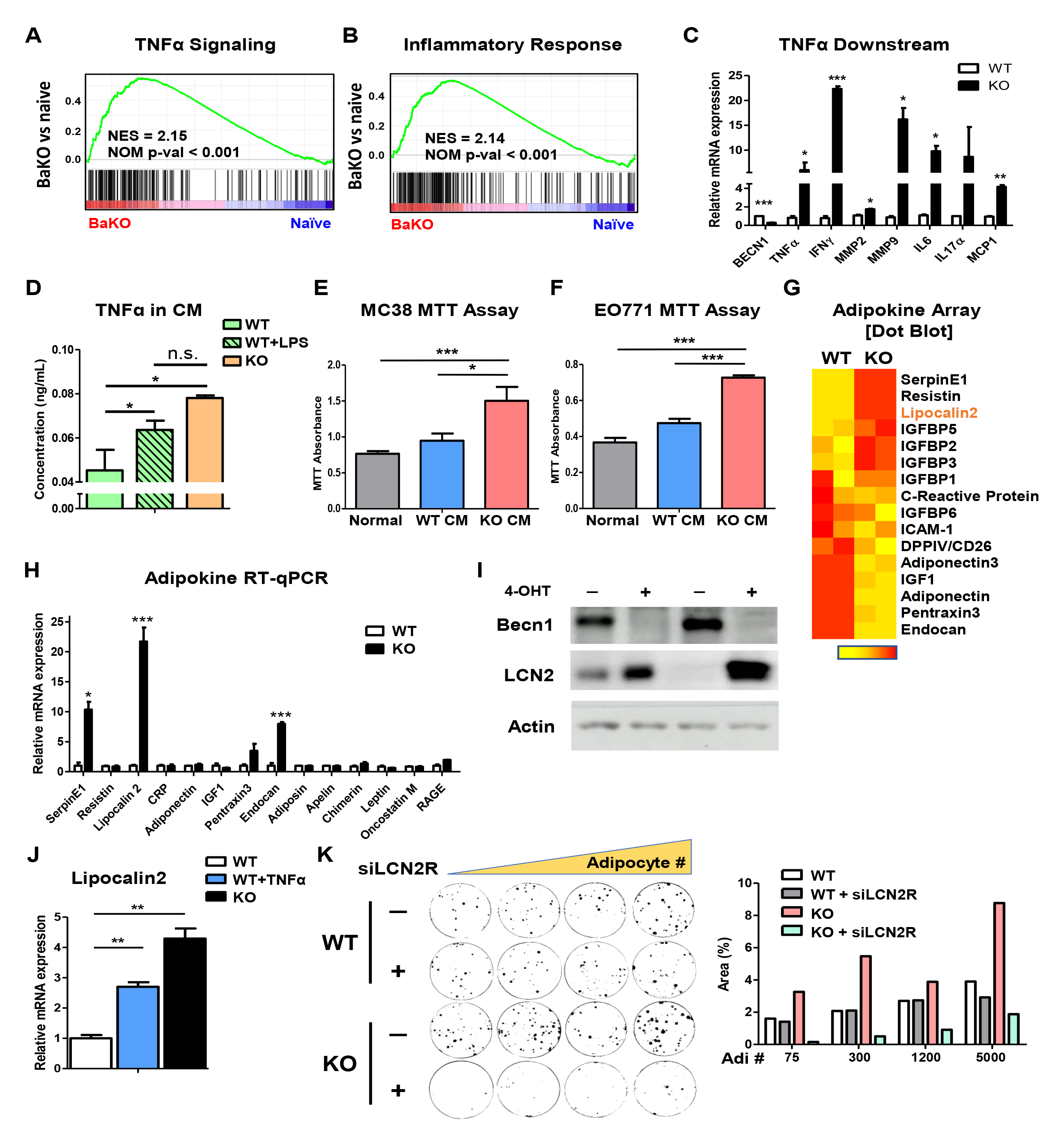
BECN1-deficient adipocytes induce inflammatory signals, TNFα and LCN2, to support tumor growth. A GSEA plot for TNFα signaling (A) and inflammatory response gene sets (B) compared between WT and BaKO iWATs. C Relative mRNA expression of TNFα downstream genes between WT and *Becn1 KO* imSVCs (n=3). D Concentration of TNFα in the CM extracted from imSVCs of indicated genotypes using ELISA (n = 3). LPS (100 ng/mL) was treated for 24 h. E Cell viability assay performed on MC-38 (E) and EO771 (F) grown in normal or conditioned media extracted from imSVCs for 72 h (n = 3). G Heatmap analysis of mouse adipokine array tested on WT and BaKO serum (n = 2, each genotype). H Relative mRNA expression of adipokines associated with tumor growth (n = 3). I Western blot analysis of protein lysates isolated from imSVCs. J Relative mRNA expression of LCN2 between imSVCs treated with or without TNFα recombinant proteins for 24 h (n = 3). K Colony assay of EO771 co-cultured with WT or BECN1 KO adipocytes. A total of 50 cancer cells were seeded with or without siLCN2R treatment for 2 days before the co-culture (co-cultured adipocyte numbers are as indicated). Area (%) was quantified using ImageJ. Data information: Two-tailed unpaired Students *t-*test (C, D, E, F, H, and J). Data are shown as mean ± SEM; \**p* ≤ 0.05, \*\**p* ≤ 0.01, \*\*\**p* ≤ 0.001.

TNFα, among other inflammatory factors, has been shown to impair adipocyte differentiation, adipogenic potential, and fat storage (Cawthorn & Sethi, 2008). Notably, adipocytes co-cultured with cancer cells display a marked increase in TNFα signaling (Fig EV3A), which can regulate the expression of various adipokines. Therefore, we tested whether secretory factors produced by BECN1-deficient adipocytes are sufficient to promote cancer cell growth. Using centrifugal filters, we concentrated the CM obtained from WT and BECN1-deficient adipocytes and transferred them to cancer cells. Treatment with CM extracted from BECN1-deficient adipocytes led to accelerated growth of both MC-38 and EO771 compared to that of CM derived from WT adipocytes (Fig. 4E and F).

Adipokines are signaling molecules secreted by adipocytes that play critical roles in regulating cellular processes, such as cell proliferation, angiogenesis, and inflammation, all of which are essential in tumor growth and metastasis (Lengyel *et al*, 2018; Rybinska *et al*, 2020; Rybinska *et al*., 2021). To further interrogate adipokine secretion and its potential role in tumor growth, we measured the levels of 38 adipokines in BaKO plasma using a mouse adipokine array kit (Fig EV3B). We then quantified the expression of each adipokine and aligned them in the order of significance (Fig. 4G) and found that BaKO plasma contained higher levels of oncogenic cytokines, such as SerpinE1, Resistin, LCN2, and insulin-like growth factor binding proteins. However, since these adipokines could be secreted from multiple organs systemically, we inspected their expression in WT and BECN1-deficient adipocytes. Among multiple adipokines, LCN2 was consistently upregulated in BECN1-deficient adipocytes (Fig. 4H and I). Further, we found that adipocyte-LCN2 could be amplified by delivering recombinant TNFα protein (Schroder *et al*, 2020), suggesting that constitutively active TNFα signaling in BaKO iWAT is responsible for LCN2 secretion (Fig. 4J; Fig EV3C).

LCN2 has been implicated in the development of metabolic disorders (Yan *et al*, 2007), and its levels are upregulated in the adipose tissue of obese patients with low-grade inflammation (Wang *et al*, 2007). Moreover, LCN2 is a pro-inflammatory signal that is involved in promoting cancer cell survival and malignancy (Yang & Moses, 2009). Consistent with these findings, we found that recombinant LCN2-protein treatment increased the proliferation of MC-38 and EO771 cancer cell lines in a dose-dependent manner (Fig EV3D and E). Next, we assessed the impact of LCN2 uptake from cancer cells co-cultivated with BECN1-deficient adipocytes. Knock-down of the LCN2-receptor (SLC22A17) in cancer cells was sufficient to attenuate the effect of *Becn1 KO* adipocytes (Fig. 4K). These results show that BECN1 depletion induces inflammatory and oncogenic adipokine secretion, especially TNFα and LCN2, to provide a pro-tumorigenic niche.

### BECN1 regulates YAP/TAZ independent of autophagy

We previously showed that the loss of adipocyte BECN1 resulted in autophagy inhibition (Jin *et al*., 2021). To determine if the autophagy-related function of BECN1 drives adipocyte transformation, we tested the effects of autophagy inhibitors, bafilomycin A and hydroxychloroquine (HCQ), on adipocytes. Although autophagy flux was clearly blocked in mature adipocytes, we did not observe any notable changes in YAP/TAZ expression or its downstream signaling (Fig. 5A and B; Fig EV3F). Additionally, while *BECN1* deletion in HEK293 induced nuclear translocation of YAP/TAZ, autophagy-related gene 7 (*ATG7*) deletion could not reproduce this effect (Fig. 5C and D). Furthermore, we employed adipocyte-specific *Atg7 KO* mice (ATG7aKO) to assess the tumor progression of MC-38 and EO771. Unlike BaKO, *Atg7* deletion was insufficient to provide a pro-tumorigenic environment (Fig. 5E and F; Fig EV3G and H). A similar result was obtained when the mice were treated with HCQ (Fig. 5G). Studies of BECN1 have highlighted its dynamic roles in regulating cellular homeostasis beyond autophagy. Its involvement in PI3K/Akt/mTOR signaling and other pathways to modulate cell growth and survival has proposed *BECN1* as a tumor suppressor gene (Hu *et al*., 2020; Liang *et al*., 1999; Qu *et al*., 2003; Wijshake *et al*, 2021; Yue *et al*., 2003). Collectively, the alteration of YAP/TAZ expression and adipocyte transformation observed in BaKO could be attributed to the autophagy-independent function of BECN1.

**Figure 5.**
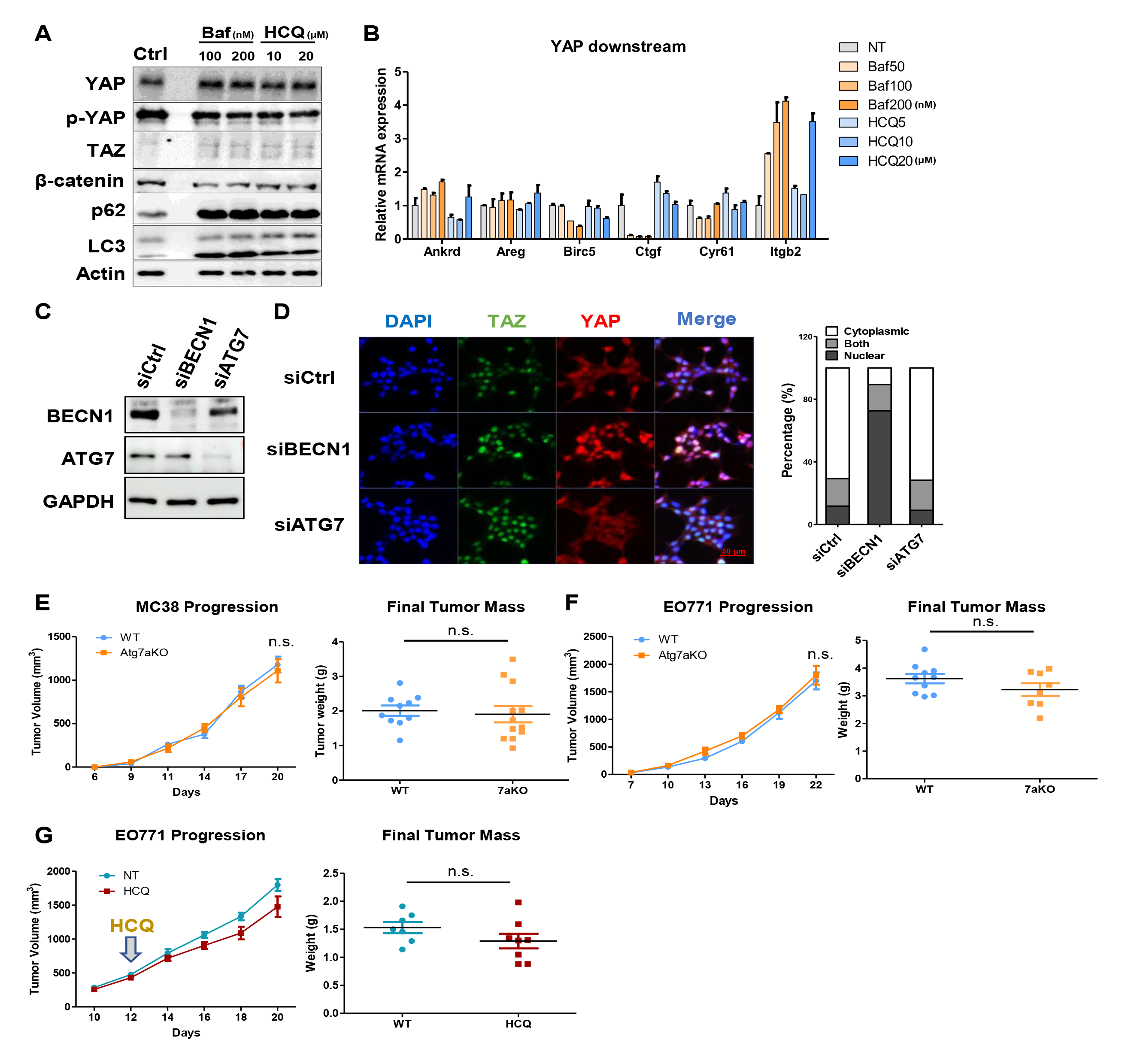
BECN1 regulates YAP/TAZ independent of autophagy. A Western blot analysis of protein lysates isolated from differentiated 3T3L1 treated with autophagy inhibitors, Bafilomycin A (Baf) and HCQ, for 48 h. B Relative mRNA expression of YAP downstream signaling genes in differentiated adipocytes (10T1/2) treated with autophagy inhibitors for 48 h. C Western blot analysis of protein lysates isolated from HEK293T cells treated with siCtrl, siBECN1, and siATG7. D Representative IF image and quantified graph of HEK293T cells stained with YAP, TAZ, and DAPI after siRNA BECN1 and ATG7 transfection. E Tumor volumes and weights measured after subcutaneous injection of MC-38 into 7- week-old WT and ATG7aKO (WT, n =10; ATG7aKO, n = 12). F Tumor volumes and weights measured after mammary fat pad injection of EO771 into 7-week-old WT and ATG7aKO (WT, n = 10; ATG7aKO, n = 8). G Tumor volumes and weights measured after mammary fat pad injection of EO771 into 7-week-old mice treated with and without HCQ (50 mg/kg, injected everyday) (NT, n = 7; HCQ, n = 8). Data information: Ordinary two-way ANOVA (E, F, and G) and two-tailed unpaired Students *t-*test (E, F, and G). Data are shown as mean ± SEM; \**p* ≤ 0.05, \*\**p* ≤ 0.01, \*\*\**p* ≤ 0.001.

### Generation of *Becn1/Yap1/Taz KO* mice to restore BaKO phenotype

The Hippo pathway is a critical signaling pathway responsible for maintaining adipocyte homeostasis (Lorthongpanich *et al*, 2019; Shen *et al*, 2022). Our data highlights that BECN1 depletion results in amplified YAP/TAZ signaling, which is associated with altered adipocyte characteristics, such as EMT and adipogenesis (Lorthongpanich *et al*., 2019; Oliver-De La Cruz *et al*, 2019; Shen *et al*., 2022). Furthermore, we have demonstrated that YAP/TAZ signaling is increased when adipocytes interact with cancer cells (Fig. 2C and I). To eliminate YAP/TAZ-mediated adipocyte transformation, we crossed BaKO with YAP/TAZ-floxed mice and generated adipocyte-specific *Becn1/Yap1/Taz* triple-KO mice (BYTaKO) (Fig. 6A; Fig EV4A). Intriguingly, although autophagy remained to be impaired (Fig EV4B), BYTaKO showed restored phenotypes in the adipose tissue and liver compared to BaKO (Fig. 6B-D; Fig EV4C). While lipodystrophy observed in BaKO was alleviated in BYTaKO, these results were only achieved by simultaneous deletion of *Yap1* and *Taz* (Fig EV4D).

**Figure 6.**
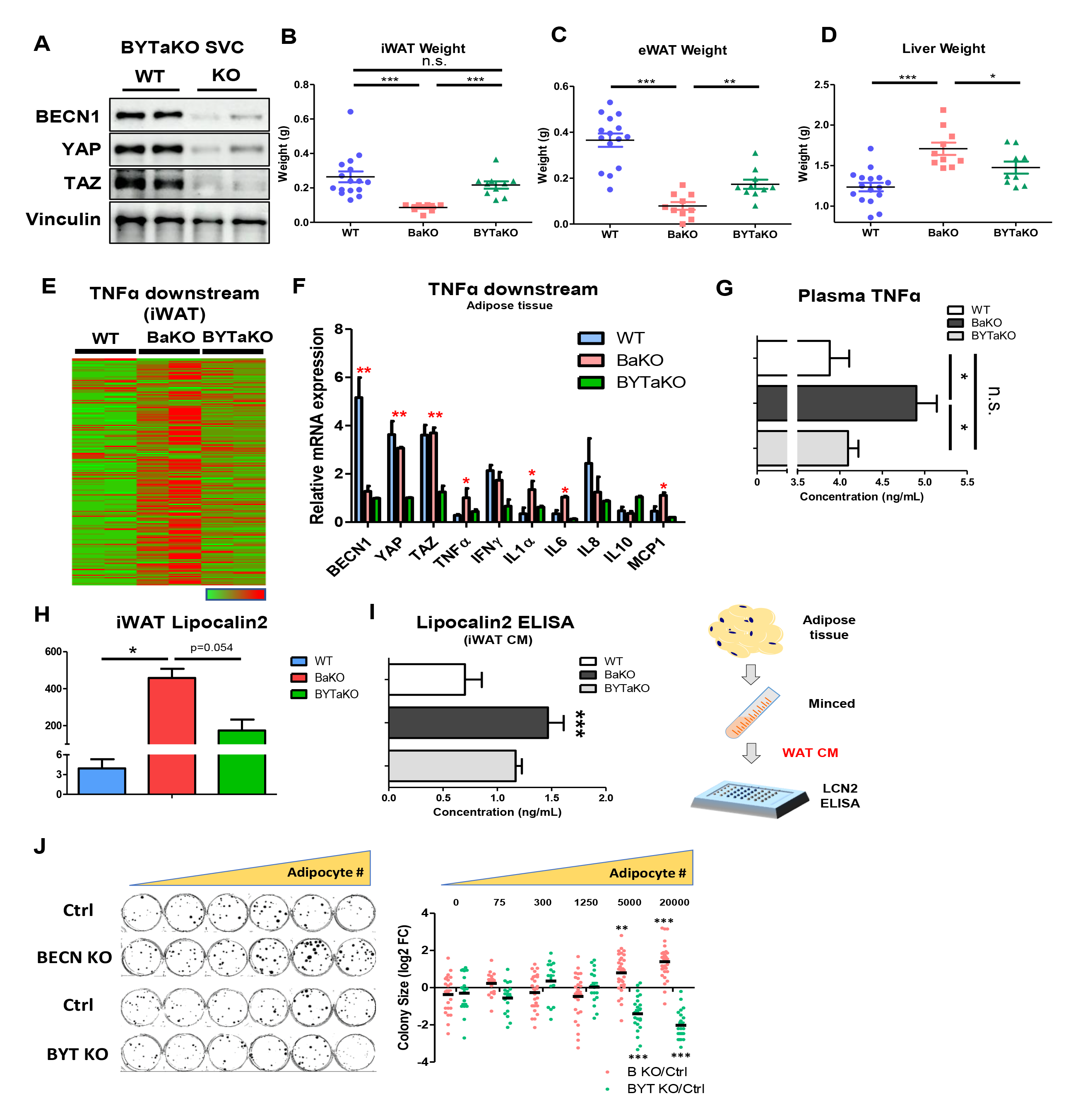
BaKO phenotypes are restored in BECN1/YAP/TAZ triple KO mice. A Western blot analysis of protein lysates isolated from fully differentiated WT and BYTaKO SVCs. B, C, and D Organ weights measured from 8-week-old WT, BaKO, and BYTaKO (WT, n = 16; BaKO, n = 10; BYTaKO, n = 10). E Heatmap analysis of TNFα downstream gene expression obtained from WT, BaKO, and BYTaKO iWAT. F Relative mRNA expression of BECN1, YAP, TAZ, and TNFα downstream genes confirmed by RT-qPCR (n = 2 mice per genotype). G Concentration of TNFα in mouse serum using ELISA (WT, n = 5; BaKO, n = 4; BYTaKO, n = 3). H Relative mRNA expression of LCN2 mouse iWAT using RT-qPCR (n = 2 mice per genotype). I Concentration of LCN2 in iWAT CM extracted from minced adipose tissue (n = 3). J Colony assay of EO771 co-cultured with doxy-inducible BECN1 and BECN1/YAP/TAZ KO adipocytes. A total of 30 cancer cells were seeded with differentiated adipocytes (adipocyte number indicated). The sizes of each EO771 colony were quantified using *ImageJ* and plotted relative to control in log2-fold change (FC). Data information: Two-tailed unpaired Students *t-*test (B, E, F, G, and H) and one-way ANOVA (D and G). Data are shown as mean ± SEM; \**p* ≤ 0.05, \**\*p* ≤ 0.01, \*\*\**p* ≤ 0.001.

Wang *et al*. demonstrated the protective function of YAP/TAZ in adipocytes under cellular stress, where they elaborate on the mechanism of TNFα-mediated YAP/TAZ nuclear translocation (Wang *et al*, 2020). BECN1-deficient adipocytes exhibited enhanced YAP/TAZ nuclear translocation followed by an activation of TNFα signaling (Fig. 4A-D; Fig EV1A-C). Accordingly, transcriptomic analyses revealed that enhanced TNFα signaling observed in BaKO was curtailed in BYTaKO iWAT (Fig. 6E and F). The plasma TNFα concentration was also reduced by the additional deletion of YAP/TAZ (Fig. 6G), implying that systemic inflammation was mitigated in BYTaKO. Thus, the healthy adipose tissue properties of BYTaKO mice led them to resist HFD-induced obesity (Fig EV4E and F). Furthermore, LCN2 expression was downregulated in BYTaKO compared to that in BaKO (Fig. 6H). We collected WAT CM to measure LCN2 levels and found that BYTaKO WAT contained lower levels of LCN2 than BaKO (Fig. 6I).

To assess whether BECN1/YAP/TAZ (BYT)-deficient adipocytes could impact tumor growth, we generated doxycycline-inducible BYT KO adipocytes and performed a colony formation assay of EO771 grown with WT, BECN1, and BYT-deficient adipocytes. When grown with BECN1-deficient adipocytes, cancer cells exhibited an enhanced proliferation rate as the number of adipocytes increased (Fig. 6J). Conversely, the opposite effect was observed when the cancer cells were grown with BYT-deficient adipocytes (Fig. 6J), suggesting that BYT-deficient adipocytes interacted with cancer cells to suppress their growth. Therefore, additional YAP/TAZ deletion in BYTaKO protected adipocytes from transformation, mitigating excessive inflammatory response and LCN2 secretion compared to BaKO.

### Inhibition of YAP/TAZ activity suppresses the tumorigenic effect of BECN1-deficient adipocytes

To validate the effect of BYT-deficient adipocytes *in-vivo*, we transplanted MC-38 and EO771 into WT, BaKO, and BYTaKO. Coherent with our transcriptomic analysis and colony formation assay, tumor growth was significantly suppressed in BYTaKO compared to that in BaKO (Fig. 7A and B). We next explored the interplay between each mouse genotype, including the peritumoral WATs. To gain a preliminary understanding of the gene expression patterns in WATs, we performed principal component analysis (PCA) on RNA-seq results of naïve and peritumoral WATs (WT, BaKO, and BYTaKO). Notably, we observed significant difference between BaKO and WT naïve WATs, whereas the BYTaKO WAT exhibited a milder phenotype. This difference was even more pronounced in peritumoral WATs, supporting the idea that BYTaKO WAT underwent a milder transformation upon cancer transplantation (Fig EV5A). As previously reported, the transformation of adipocytes near cancer cells is associated with the downregulation of mature adipocyte markers (Bochet *et al*., 2013; Zhu *et al*, 2022a; Zoico *et al*., 2016) (Fig. 2B, G, and H). Accordingly, the expression of adipogenesis-and fatty acid (FA) metabolism-associated gene sets indicated that the adipocytes in peritumoral WAT of all genotypes had lost adipogenic potential and mature adipocyte properties (Fig. 7C). However, BaKO WATs exhibited significantly lower levels of adipogenesis-and FA metabolism-associated genes, whereas BYTaKO WATs retained relatively higher levels of these genes (Fig. 7C).

**Figure 7.**
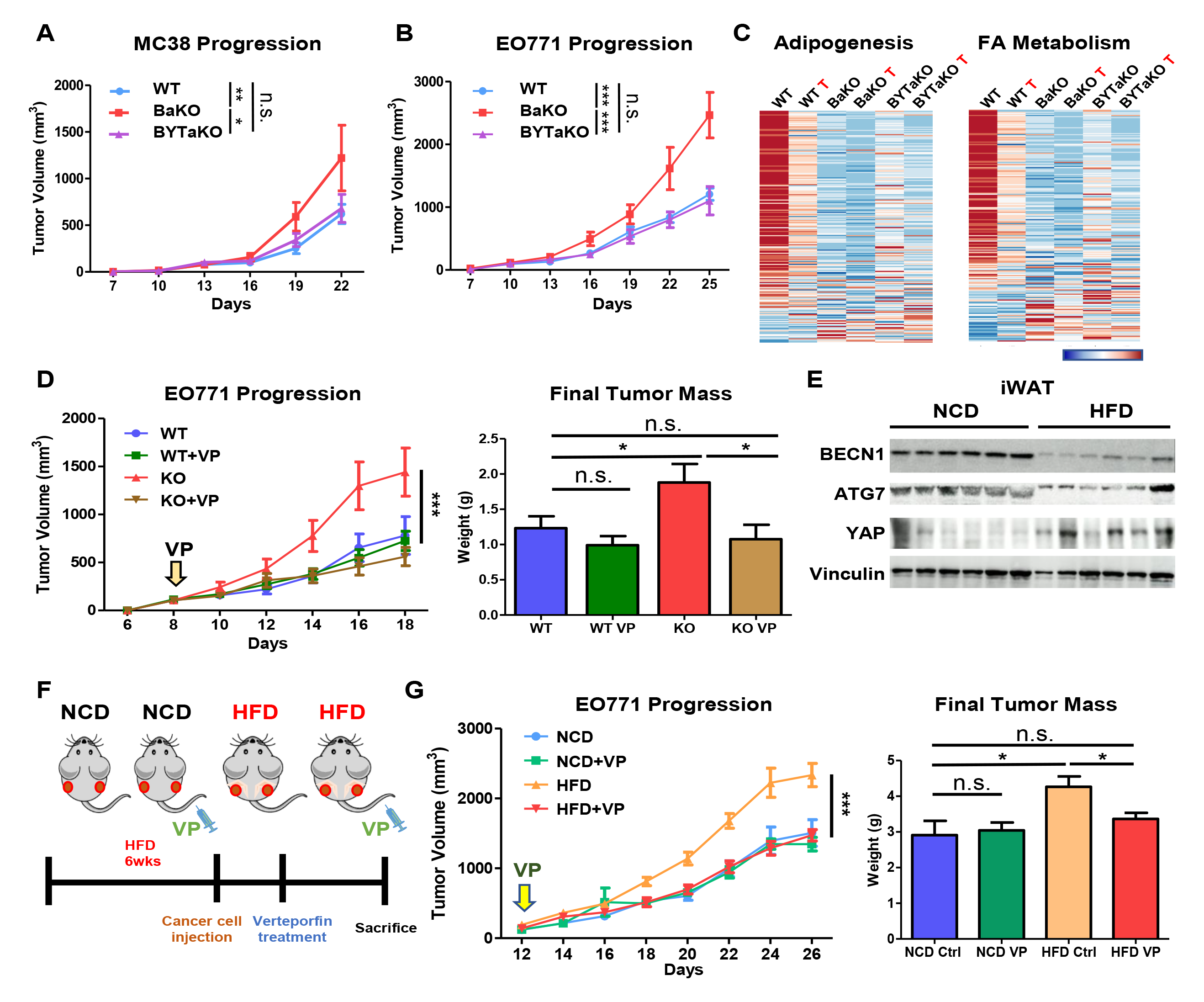
Inhibition of YAP/TAZ activity suppresses the tumorigenic effect of BECN1-deficient adipocytes. A and B Tumor volumes measured after MC-38 (A) and EO771 (B) injected into 7-week-old WT, BaKO, and BYTaKO. C Heatmap analysis of adipogenesis and FA metabolism gene sets between WT, BaKO, and BYTaKO iWATs and their peritumoral WATs (labeled with red letter ‘T’). D Tumor volumes and weights after mammary fat pad injection of EO771 into 7- week-old mice treated with or without VP (30 mg/kg, injected every other day). VP treatment was initiated when tumors reached a volume of 150 mm^3^, and the mice were sacrificed on day 22 following injection (WT, n = 8; WT VP, n = 8; KO, n = 6; KO VP, n = 6). E Western blot analysis of protein lysates isolated from iWAT of NCD and HFD-fed mice. Mice were fed with HFD chow for 12 weeks. F Schematic timeline of NCD and HFD-fed mice treated with or without VP. G Tumor volumes and weights after mammary fat pad injection of EO771 into NCD or HFD-fed mice. VP treatment (30 mg/kg, injected every other day) was initiated when tumors reached a volume of 150 mm^3^, and the mice were sacrificed on day 26 following the injection. Data information: Ordinary two-way ANOVA (A, B, D, and G) and two-tailed unpaired students t-test (D and G). Data are shown as mean ± SEM; \**p* ≤ 0.05, \*\**p* ≤ 0.01, \*\*\**p* ≤ 0.001.

Several studies have demonstrated the anti-tumor activity of verteporfin (VP) through suppression of YAP/TAZ signaling (Feng *et al*, 2016; Wang *et al*, 2016). To complement our genetic mouse model (BYTaKO), we employed VP to inhibit YAP/TAZ activity in adipose tissue (Fig EV5B and C). To avoid the direct effect of VP on cancer cells, we administered a lower dose of VP that did not affect WT tumor growth (Fig. 7D). However, VP treatment led to a regression of tumor in BaKO, similar to that observed in BYTaKO, highlighting the potential impact of VP on tumors growing in adipose-rich environments (Fig. 7D). We also hypothesized that VP could induce tumor regression in mice with metabolic dysregulation. Diet-induced obesity mice develop various metabolic disorders involving inflammatory adipose tissue, and we examined the relationship between BECN1 and YAP expression in the iWAT of HFD-fed mice. Interestingly, while BECN1 levels were decreased, YAP levels were increased in iWATs but not in eWATs, indicating the formation of a malignant TME (Fig. 7E; Fig EV5D). Therefore, we evaluated tumor progression in HFD-fed mice and found that EO771 grew more rapidly (Fig. 7F). However, VP treatment led to tumor regression only in HFD-fed mice (Fig. 7G; Fig EV5E-G). These data indicate that BECN1 and YAP levels serve as indicators of adipose tissue health and their potential to provide a pro-tumorigenic environment. Consequently, our findings advocate the use of YAP/TAZ inhibitors such as VP in patients with metabolic dysregulation to enhance its anti-tumor activity (Fig. 8A).

**Figure 8.**
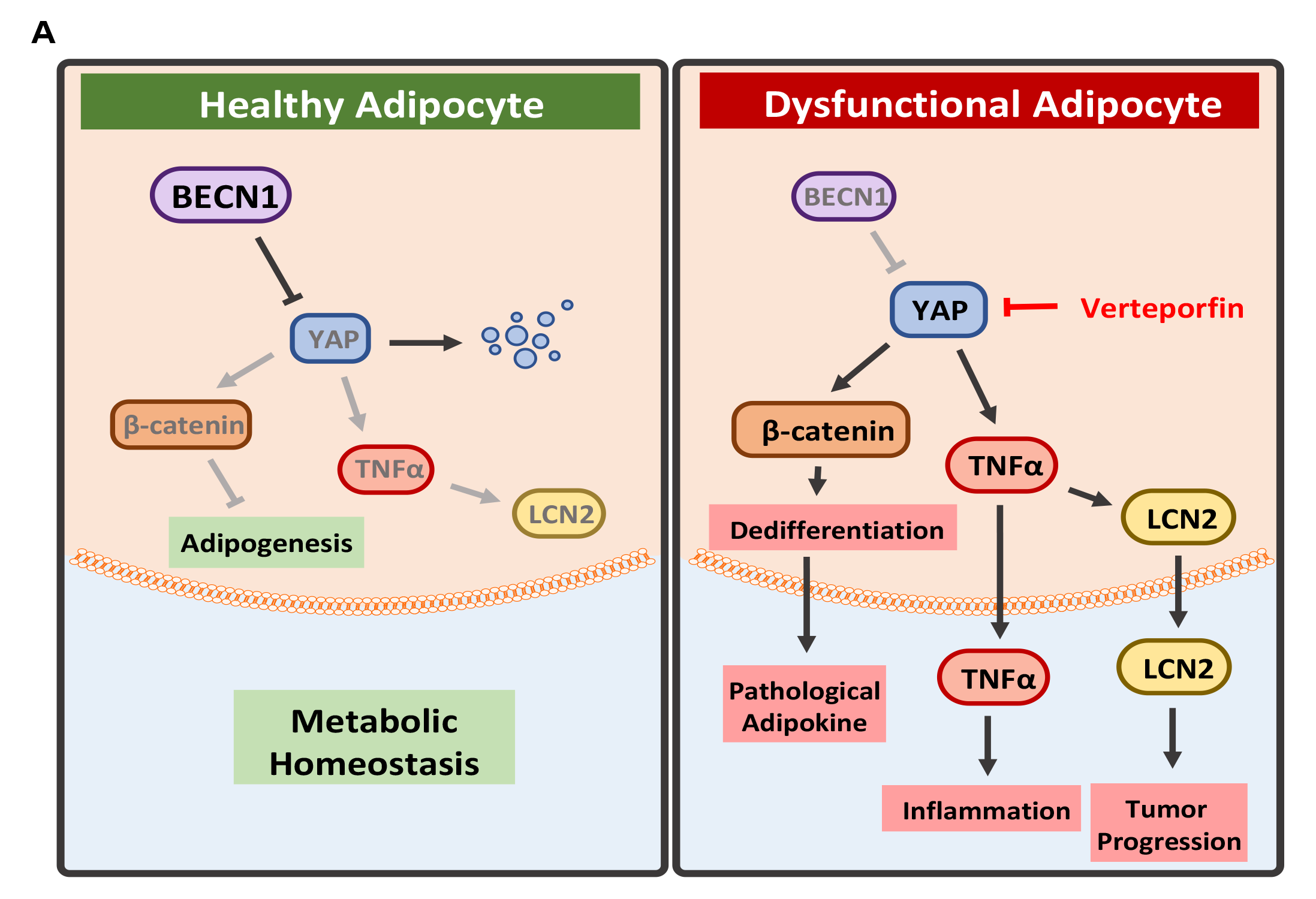
Illustration of the role of BECN1 in retaining adipocyte homeostasis. A Schematic illustration of signaling mechanism identified in this study.

## Discussion

Numerous studies have shown that autophagy-associated proteins may have functions that extend beyond autophagy. For example, the tumor suppressive role of UVRAG and BECN1 could be attributed to the trafficking function of BECN1/UVRAG in initiating autophagy (Wijshake *et al*., 2021), which regulates the cell surface localization of E-cadherin. This, in turn, prevents tumor progression *via* contact inhibition of proliferation through Hippo signaling, suppression of β-catenin, and loss of mesenchymal phenotype (Wijshake *et al*., 2021). Thus, these properties of Beclin1 could contribute to the adipocyte transformation observed in BECN1-deficient adipocytes and BaKO. In addition to its trafficking function, BECN1 has been found to directly interact with various proteins, including Hippo-associated proteins, such as LATS1 and MST1 (Maejima *et al*, 2013; Tang *et al*, 2019). Therefore, the mechanism by which BECN1 depletion leads to dynamic alterations in adipocyte properties, including YAP/TAZ signaling, extends beyond autophagy inhibition. Furthermore, given that BECN1 has a tumor suppressive function in multiple cancer types, a more comprehensive understanding of Beclin1 outside of autophagy is necessary.

Previous research has extensively investigated the role of the Hippo pathway in cancer cells, but our study highlights the importance in adipocytes as a component of the TME. Specifically, TAZ has been identified as a repressor of PPARɣ, and TAZ-deficient adipocytes have exhibited improved health (El Ouarrat *et al*, 2020). Additionally, research on TAZ in adipocyte-TME has manifested its role in regulating resistin secretion, promoting tumorigenesis in triple-negative breast cancers (Gao *et al*, 2020). Our study revealed the significance of YAP/TAZ in adipocyte differentiation and homeostasis; aberrant expression of these proteins could mediate adipocyte transformation, as observed in cancer-co-cultured adipocytes. This may be due to the YAP/TAZ function to modulate cellular plasticity and stemness, which are the critical determinants of cellular fate in multiple TME compartments (Zanconato *et al*, 2019). Thus, YAP/TAZ inhibitors, such as VP, may be particularly potent as they can target both intrinsic and extrinsic factors of cancer cells, leveraging the unique properties of YAP/TAZ for a dual therapeutic benefit.

Several studies have suggested a link between cancer progression and measures such as body mass index or waist circumference (Moore *et al*, 2004; Recalde *et al*, 2021). However, these measures cannot fully capture the complexity of the TME. Our BaKO mouse model suggests that a loss of adipose tissue mass can also contribute to the development of an inflammatory TME. Interestingly, the adipocytes in BaKO WAT were not hypertrophic or hyperplastic, but still provided a pro-tumorigenic niche in the TME. Furthermore, our study found that certain diet-induced obese mice had increased levels of BECN1 and decreased levels of YAP/TAZ in their iWATs, indicating adipocyte transformation and a malignant TME. These expression patterns highlight the importance of assessing the status of adipose tissue to predict the formation of a malignant TME. Therefore, evaluating distinct expression profiles could be a more informative approach to assess adipose tissue health and predict malignant TME formation.

It is broadly accepted that CAAs undergo dedifferentiation, a complex process characterized by the morphological changes, downregulation of mature adipocyte markers, loss of lipid contents, and acquisition of mesenchymal stem cell phenotype (Na *et al*, 2023; Song & Kuang, 2019). Wnt and Notch signaling have previously been identified as mediators of adipocyte dedifferentiation for CAA generation (Bi *et al*, 2016; Zoico *et al*., 2016). However, we aimed to identify the essential attributes of adipocyte transformation and its drivers in the context of TME. Through transcriptomic analysis and co-cultivation with cancer cells, we identified the unique features of CAAs, including an excessive pro-inflammatory response and depletion of adipogenic potential. Based on these observations, we propose the term “trans-differentiation” to describe the process of adipocyte transformation in TME.

Our study demonstrated that depletion of BECN1 in adipocytes promoted YAP/TAZ signaling, leading to adipocyte transformation with similar features to CAAs. This transformation provided a pro-tumorigenic niche in the TME, enhancing breast and colon cancer progression. However, we also found that adipocyte transformation was reversible, as genetic or chemical inhibition of YAP/TAZ signaling led to tumor growth regression (Fig. 8A). These findings highlighted the critical role of adipocyte quality in determining TME malignancy, particularly in adipose-rich environments. Therefore, therapeutic interventions that improve adipocyte homeostasis should be given greater consideration in addressing malignant TME formation. Furthermore, our findings provide insights into a potential dual therapeutic strategy for targeting adipocyte transformation and tumors, which could have broad applications in treating various pathological conditions.

## Materials and Methods

### Animals

Adipocyte specific *BECN1* deficient mice (BaKO) were generated as described previously (Jin *et al*., 2021). Mice with floxed alleles of *Yap1* (Yap1^tm1a(KOMP)Mbp^), *Taz* (Wwtr1^tm1.2Eno^), and *Atg7* (Atg7^tm1Tchi^ RBRC02759; RIKEN) were obtained from the Knockout Mouse Project (KOMP) Repository (KOMP, Davis, California, USA). By crossing BaKO with YAP^flox/flox^ (Yap1^tm1a(KOMP)Mbp^) and TAZ^flox/flox^ (Wwtr1^tm1.2Eno^), adipocyte-specific *BECN1/YAP/TAZ* KO mice (BYTaKO) were generated. Similarly, adipocyte specific *Atg7* deficient mice were generated by crossing Adipoq-Cre (JAX: 010803) with ATG7^flox/flox^ (Atg7^tm1Tchi^). The excised alleles were validated using PCR analysis of genomic DNA extracted from the tail tips of mice. The genotyping primer sequences are listed in Appendix Table S1.

Furthermore, Adipoq-cre, BECN1^flox/flox^, MMTV-PyMT (PyBA) were generated by crossing BaKO with MMTV-PyMT (The Jackson Laboratory, Bar Harbor, ME, USA). The integrated MMTV-PyMT sequence was validated using PCR analysis of genomic DNA extracted from tail tips of mice. Mice received either a normal chow diet (NCD) or high-fat diet (HFD) (consisting of 60 kcal %), in addition to water, *ad libitum*, for indicated times. To carry out the experiments, immunocompromised N2G (NOD; Prkdc^em1Gmcr^; Il2rg^em1Gmcr^) mice were obtained from Gemcro Corp (Seoul, Republic of Korea). These mice were generated from NOD with entire exons of PRKDC and IL2RG deleted.

All animal care and experiments were performed in accordance with the guidelines of the Korean Food and Drug Administration and approved by the Institutional Animal Care and Use Committees of the Laboratory Animal Research Center at Yonsei University (permit number IACUC-A-202208-1520- 01). Female mice were used for breast cancer (EO771, 4T1, MMTV-PyMT) related experiments, while male mice were used for colon cancer (MC38) related experiments. The age of each mouse cohort is indicated, and their weights ranged from 20 to 30 g. Mice were maintained in the specific pathogen-free facility at Yonsei Laboratory Animal Research Center.

### Adipose tissue fraction

Adipose tissues were isolated and fractionated as described previously (Jin *et al*., 2021). WATs were isolated, dissected out, chopped, and incubated in collagenase buffer for 20 min at 37 °C with shaking. Cell suspensions were centrifuged to separate adipocyte (supernatant) and stromal vascular fractions (pellet). Peritumoral adipose tissues were obtained two weeks after the injection of 1 × 10^6^ EO771 cells at the region within 5 mm of the primary tumor.

### Cell cultures and adipocyte differentiation

*In vitro* cultures were performed as described previously (Jin *et al*., 2021). Immortalization of stromal vascular cells (SVC) and induction of adipocyte differentiation were conducted as previously described (Jin *et al*., 2021). For co-culture experiments, differentiated adipocytes and EO771 (ATCC) were cultured together in 60 pi (SPL #209260) or 6-well (SPL #37006) co-culture chambers. Colony formation assays were performed by mixing various numbers of differentiated adipocytes (incubated for 2 days in differentiation media and then 4 days with DMEM (Gibco), 1 μM insulin (I9278, Sigma-Aldrich), 10% FBS (Gibco), and 1% pen/strep (Gibco) with the EO771 breast cancer cell line (Jin *et al*., 2021). After staining the cells with crystal violet (548-62-9, Sigma-Aldrich), colony sizes were measured using ImageJ 1.53t (Bethesda, MD, USA). To prepare the conditioned media, differentiated adipocytes were incubated for 48 h in a medium consisting of DMEM, 1% FBS, and 1% pen/strep. The conditioned media (CM) was then 5-fold concentrated using a centrifugal filter (Merk, C7715) and mixed with DMEM, 3% FBS, and 1% pen/strep in a 1:1 ratio before treatment. Cell viability was measured using the MTT assay (M6494; Invitrogen).

### Syngeneic and xenograft models

MC-38 and EO771 cells were injected into the subcutaneous or mammary fat pads of WT, BaKO, or BYTaKO mice, and tumor sizes were periodically measured using the formula:

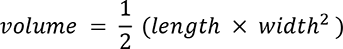

The mice were euthanized by cervical dislocation, and the tumor weights were measured after resection. To study the effect of adipocyte differentiation on tumor growth, immortalized SVCs (imSVCs) pre-treated with differentiation media were mixed with 4T1 cells and injected subcutaneously into N2G mice. To deplete BECN1, mice were treated with 4-hydroxytamoxifen (4- OHT) for two days.

### Retroviral shRNA design and transduction

The lentivirus inducing shBECN1 was generated through transfection with psPAX2, pmD2G, and pLV-H1TetO-GFP-Puro using Lipofectamine 3000 (Thermo Fisher # L3000001) following the manufacturer’s instructions. For retroviral transduction of 10T1/2 (ATCC), polybrene (Merk #TR-1003- G) was used. The transduced cells were selected by incubation in medium containing DMEM, 1 μM insulin, 10% FBS, 1% pen/strep, and 2 µg/mL puromycin. The shRNA oligonucleotides targeting mouse BECN1, YAP, and TAZ were designed using the web service (https://biosettia.com/support/shrna-designer/) and their sequences are listed in Appendix Table S2.

### RNA interference-mediated knockdown

The transfection of siATG7 (SC-29797; Santa Cruz), siBECN1 (SC-41447; Santa Cruz), and siLCN2R was carried out using Lipofectamine 3000 (Thermo Fisher # L3000001), following the manufacturer’s instructions. The siLCN2R was designed using the web service (https://biosettia.com/support/shrna-designer/) and synthesized by GenePharma. The siLCN2R mixture was prepared by combining 2.5 ng/μL of each siRNA to yield a final concentration of 10 ng/μL. The sequences for siLCN2R are provided in Appendix Table S3.

### Glucose tolerance test

Glucose tolerance was assessed by administering a 20% glucose injection (1 g/kg body weight), as described in a previous study (Jin *et al*., 2021).

### Drug uptake

Drug uptake was assessed by administering 10 mg/mL Verteporfin (129497-78-5, USP) dissolved in 5% dimethyl sulfoxide (DMSO) corn oil (30 mg/kg or 50 mg/kg) *via* intraperitoneal injection in WT and BaKO mice.

### Quantification of free fatty acids, adipokines, and cytokines

The concentration of free fatty acids in the adipocyte-CM was quantified using an ELISA assay kit (ab65341; Abcam). The levels of adipokines in the plasma of WT and BaKO mice were measured using the Mouse Adipokine Array Kit (ARY013; R&D Systems) and LEGENDplex (740929; Biolegend). Blood was collected from the retroorbital sinus using a capillary tube.

### Western blotting and antibodies

Western blotting was performed as previously described (Jin *et al*., 2021) using primary antibodies against BECN1 (Cell Signaling Technology #3738 and Santa Cruz #sc-48341), FASN (#3180; Cell Signaling Technology), PPARγ (#2435; Cell Signaling Technology), PLIN1 (#9349; Cell Signaling Technology), LC3 (Sigma-Aldrich #L8918), p62 (Abnova #H00008878-M01), actin (Santa Cruz Biotechnology #sc-47778), GAPDH (Santa Cruz Biotechnology #sc-32233), tubulin (Santa Cruz #sc-48341), β-catenin (Santa Cruz #sc-7963), YAP/TAZ (Santa Cruz #sc-101199), YAP (#14074; Cell Signaling Technology), TAZ (#4883; Cell Signaling Technology), 12hosphor-YAP (#13008; Cell Signaling Technology), Lipocalin2 (LCN2; R&D systems #AF1857), HSL (#4107; Cell Signaling Technology), and phosphor-HSL (#4137, #4139, #45804; Cell Signaling Technology).

### Immunofluorescence (IF) imaging microscopy

Before cell seeding, a cover glass (Marienfield) was coated with a 1% (w/v) gelatin solution in PBS and incubated for 30 min at 37 ℃. After washing the cells with PBS, they were fixed with 10% formalin in PBS for 15 min at room temperature (RT). Following three washes with PBS, the cells were permeabilized with 0.5% Triton X-100 in PBS for 10 min at RT. After three additional washes with 0.1% Triton X-100 in PBS, the cells were blocked with 5% BSA and 0.5% Triton X-100 in PBS for 30 min at RT. The cover glass was placed on parafilm inside a humid chamber and incubated for 2 h at RT with the following primary antibodies, which were diluted to 1:100 in 5% BSA and 0.5% Triton X-100 in PBS: anti-BECN1 (#3738; Cell Signaling Technology and sc-48341; Santa Cruz Biotechnology), anti-PLIN1 (#9349; Cell Signaling Technology), anti-β-catenin (sc-7963; Santa Cruz Biotechnology), anti-YAP/TAZ (sc-101199; Santa Cruz Biotechnology), anti-YAP (#14074; Cell Signaling Technology), and anti-TAZ (#4883; Cell Signaling Technology). After washing the cells three times with 0.1% Triton X-100 in PBS, 5 min per wash, they were incubated with secondary antibodies conjugated with fluorophores (A32744 and A21206; Invitrogen) diluted to 1:100 in PBS with 0.1% Triton X-100. Following three additional washes with 0.1% Triton X-100 in PBS for 5 min each, the cells were mounted on glass slides using Fluoroshield with 4’,6-diamidino-2-phenylindole (DAPI) (F6057; Sigma-Aldrich). Conventional fluorescent imaging was performed using the Axio Oberver.Z1/7 (Carl Zeiss), while confocal images were captured using LSM980 (Carl Zeiss).

### Histology

Tissue sampling for histology was performed as previously described (Jin *et al*., 2021).

### RNA extraction, reverse transcription quantitative real-time PCR (RT-qPCR), and RNA sequencing (RNA-seq)

RNA extraction and reverse transcription were performed according to the established protocol (Jin *et al*., 2021). For quantitative PCR analysis, primers were designed and listed in Appendix Table S4. The data were normalized to the expression levels of housekeeping genes, *GAPDH* and *β-actin*. RNA-seq was carried out by Geninus following the protocol outlined in Appendix Table S5. The RNA integrity number of the samples used for RNA-seq was above 9. The Gene Set Enrichment Analysis (GSEA) and fold change analysis were calculated based on transcripts per million values.

### Statistical analysis

The data are presented as mean ± SEM. Statistical analysis was performed using GraphPad Prism 5 software (Graph-Pad Software, Inc.). To compare two groups, the unpaired Student *t*-test was used. For comparisons of more than two groups, a two-way ANOVA was employed. Repeated-measures ANOVA was used to analyze body weight and GTT data. The Bonferroni post hoc test was used to determine significant differences after ANOVA. Statistical significance was considered at *p <* 0.05.

## Acknowledgments

This study was funded by National Research Foundation of Republic of Korea: Ministry of Science and ICT (2020R1A4A1019063), National Cancer Center (HA22C0147), and Ministry of Education (2022R1A6A3A13071199). We acknowledge the support from Brain Korea 21 (BK21) PLUS program and Yonsei University.

## Author’s Contributions

**Y. Song:** Data Curation, validation, investigation, visualization, methodology, writing-original draft and editing. **H. Na:** Data Curation, validation, investigation, visualization, methodology, writing-original draft. **S.E. Lee:** Software, methodology. **J. Moon:** Data curation. **T.W. Nam:** Data curation. **Y.M. Kim:** Data curation. **Y. Ji:** Methodology, writing-review. **Y. Jin**: Methodology, investigation. **J.H. Park:** Data curation. **S.C. Cho:** Data Curation. **D. Hwang:** Bioinformatics, conceptualization. **J.B. Kim:** Conceptualization, writing-review. **S.-J. Ha:** Conceptualization, resources. **H.W. Park:** Conceptualization, resources. **H.-W. Lee:** Conceptualization, resources, supervision, investigation, writing-review, editing, project administration, and funding acquisition.

## Data availability

All data were generated by the authors and included in the article. RNA-seq data used in this study are available upon request. To obtain the data, please contact Han-Woong Lee.

## Expanded View Figure Legends

**Figure EV1 - BECN1 depletion leads to adipocyte transformation.**

A Representative IF image of mature adipocytes. Adipocytes stained with YAP/TAZ (red), PLIN1 (green), and DAPI (blue); white arrowheads indicate fully differentiated adipocytes (PLIN1 positive) with less nuclear YAP/TAZ; yellow arrowheads indicate undifferentiated adipocytes (PLIN negative) with nuclear YAP/TAZ.

B Representative IF image of doxy-inducible *Becn1 KO* adipocytes stained with BODIPY and YAP/TAZ.

C Representative IF image of doxy-inducible *Becn1 KO* adipocytes stained with YAP/TAZ (C) and β- catenin (D) with lower magnification

E Representative IF image of imSVCs indicating nuclear translocation of YAP/TAZ and its quantification.

F FFA concentration in media extracted from WT and *Becn1 KO* imSVCs

G Colony assay of 50 EO771 cells with adipocytes (numbers as indicated). The sizes of each EO771 colony were quantified using *ImageJ*.

H Live image of adipocytes co-cultured with EO771 cells for the time indicated in a trans-well plate.

**Figure EV2 - Comparison between peritumoral and BaKO WATs reveals similar traits.**

A Venn diagram depicting individual genes that were differentially expressed in BaKO and peritumoral WATs compared to naïve adipose tissue; *p* < 0.05.

B List of differentially expressed GSEA hallmark gene sets of peritumoral and naïve adipose tissue ranked by normalized enrichment score (NES). Gene sets that also appear to be differentially expressed in BaKO WAT are colored in red and blue.

C GSEA plots for epithelial mesenchymal transition gene set compared between naïve, peritumoral, and BaKO WAT.

D List of differentially expressed GSEA ‘C6 oncogenic signature’ gene sets of BaKO and naïve adipose tissue ranked by NES. Gene sets associated β-catenin and YAP are colored in blue.

E Heatmap analysis of YAP conserved signature between WT and BaKO iWATs.

F Western blotting analysis of protein lysates isolated from BaKO iWAT.

**Figure EV3 - TNFα and LCN2 are crucial for EO771 and MC-38 progression.**

A Relative mRNA expression of TNFα downstream from control adipocytes and cancer co-cultured adipocytes (n = 3).

B Mouse adipokine array performed in dot blot. Samples were extracted from mouse plasma.

C Relative mRNA expression of TNFα downstream and LCN2 after treating adipocytes with or without TNFα recombinant protein (n = 3).

D Viable cell counting of EO771 (D) and MC-38 (E) treated with LCN2 recombinant protein. Cancer cell numbers were counted as they were sub-cultured every 36 h (n = 3).

F Western blotting analysis of protein lysates isolated from differentiated adipocytes treated with autophagy inhibitors, Bafilomycin A (Baf) and HCQ, for 48 h.

G Representative image of primary tumor resected from Fig. 5E (G) and Fig. 5F (H).

**Figure EV4 - Generation of BYTaKO and the beneficiary phenotypes.**

A Mouse breeding strategy to generate BYTaKO.

B Western blot analysis of protein isolated from WT, BaKO, and BYTaKO iWATs.

C H&E staining of tissue section prepared from WT, BaKO, and BYTaKO liver.

D Body and organ weights measured from WT, BTaKO, and BYaKO mice.

E Glucose tolerance test performed on WT and BYTaKO mice after feeding with NCD or HFD for six weeks (WT, n = 4, BYTaKO n = 6, WT HFD n = 6, BYTaKO HFD n = 5).

F Body and organ weights measured from WT and BYTaKO after feeding with NCD or HFD for six weeks.

**Figure EV5 - Deactivation of adipocyte YAP/TAZ represses malignant TME formation.**

A Principal component analysis performed on RNA-seq result from WT, BaKO, and BYTaKO WATs. Peritumoral adipose tissues were extracted two weeks after MC-38 injection (n = 2).

B Tumor volumes and weights after mammary fat pad injection of EO771 into WT mice. VP treatment (50 mg/kg, injected everyday) was initiated when tumors reached a volume of 150 mm^3^, and the mice were sacrificed on day 22 following the injection.

C Relative mRNA expression of YAP downstream in WT and *Becn1* KO imSVCs treated with or without VP (20 nM) for 24 h.

D Western blot analysis of protein lysates isolated from eWAT of NCD and HFD fed mice.

E Tumor volumes and weights after subcutaneous injection of MC-38 into NCD and HFD-fed mice. VP treatment (30 mg/kg) was initiated when tumors reached a volume of 300 mm^3^, and the mice were sacrificed on day 22 following the injection.

F Image of tumors resected from Fig. 7F (F) and Supplementary Fig. S6E (G).

---

**Appendix Figure1.**
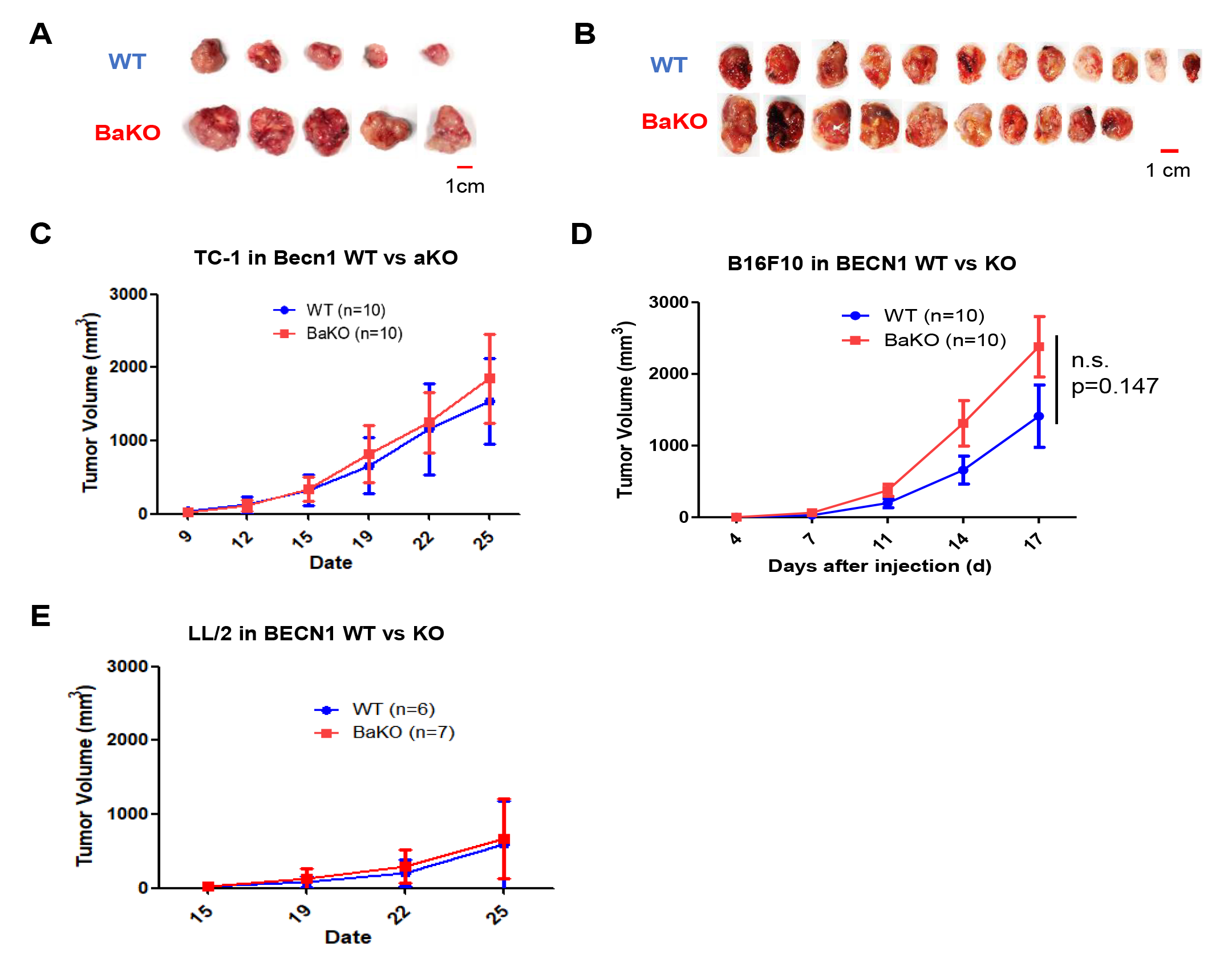
Cancer models not responsive to BECN1 deficient adipocytes. (A,B) Representative image of resected tumor presented in **Fig. 3A and B**. (C) Growth kinetics after subcutaneous injection of TC1, (D) B16F10, and (E) LL/2 into 8-week-old WT and BaKO.

